# MACanalyzeR: scRNA-seq Analysis Tool Reveals PPARγHI Lipid-Associated Macrophages Facilitate Thermogenic Expansion in BAT

**DOI:** 10.1101/2024.07.29.605588

**Authors:** Andrea Ninni, Fabio Zaccaria, Luca Verteramo, Francesca Sciarretta, Loreana Sanches Silveira, José Cesar Rosa-Neto, Paolo Grumati, Giuseppe Rizzo, Clement Cochain, Jesse W. Williams, Stoyan Ivanov, Beiyan Zhou, Katia Aquilano, Daniele Lettieri-Barbato

**Affiliations:** Department of Biology, University of Rome Tor Vergata, Rome, Italy; PhD Program in Evolutionary Biology and Ecology, Department of Biology, University of Rome Tor Vergata, Italy; IRCCS Santa Lucia Foundation, Rome, Italy; Immunometabolism Research Group, Department of Cell Biology and Development, Institute of Biomedical Sciences, University of São Paulo (ICB1-USP), São Paulo 05508-000, Brazil; Telethon Institute of Genetics and Medicine, Pozzuoli, Italy; Department of Clinical Medicine and Surgery, University Federico II, Naples, Italy; Comprehensive Heart Failure Center, University Hospital Würzburg, Am Schwarzenberg 15, A15, 97078 Würzburg, Germany; Institute of Experimental Biomedicine, University Hospital Würzburg, Josef-Schneider-Str. 2, D16, 97080 Würzburg, Germany; Center for Immunology, University of Minnesota, Minneapolis; Department of Integrative Biology and Physiology, University of Minnesota, Minneapolis; Université Côte d’Azur, CNRS, LP2M, Nice, France; Department of Immunology, School of Medicine, University of Connecticut, Farmington, Connecticut, USA; IRCCS Fondazione Bietti, Rome, Italy

**Keywords:** Macrophages, scRNAseq, Immunometabolism, Thermogenesis, Adipose Tissue, Mitochondria, Machine Learning

## Abstract

Macrophages in brown adipose tissue (BAT) play a complex role in regulating its activity. However, the role of macrophages in regulating BAT activation/deactivation has not yet been comprehensively characterized. To elucidate this, we developed MACanalyzeR, a scRNAseq-based tool specifically designed to explore the macrophage features at molecular and metabolic level. MACanalyzeR was applied in scRNA-seq datasets obtained from BAT with thermogenic loss (db/db mice) and activation (High Fat Diet, HFD). Our computational approach revealed that macrophages accumulating in BAT upon these conditions resemble lipid-associated macrophages (LAMs) with foaming-like features. BAT LAMs also show a significant enrichment of genes associated with mitochondria and lysosomes. Interestingly, LAMs identified in BAT from HFD mice positively correlate with thermogenic genes and exhibit an enrichment in PPAR_γ_ signaling pathway, with an activated mitochondrial metabolism. Cell dynamic strategy, revealed that LAM with high *Pparg* expression levels (*Pparg*^HIGH^) progressively accumulate during skeletal muscle regeneration, suggesting a potential role for this LAM subcluster in maintaining tissue homeostasis. Our findings suggest *Pparg*^HIGH^ LAMs as a subclass of macrophages potentially contributing in preserving tissue homeostasis associated with high energy demand conditions such as thermogenic and regenerative stimuli.

## INTRODUCTION

Single-cell RNA sequencing (scRNAseq) is a method used to study gene expression at the single-cell level. This is in contrast to bulk RNA sequencing, which provides information only on the average expression of the predominantly represented cell population in a heterogeneous cellular system such as that of a tissue. Compelling evidence highlights the importance of brown adipose tissue (BAT) macrophages in governing adaptive thermogenesis (Villarroya et al., 2018). Recent findings have reported that macrophages participate in BAT physiology at different levels, including release of pro-thermogenic cytokines (Cereijo et al., 2018), controlling the mitochondrial quality by transmitophagy (Aquilano et al., 2023; Rosina et al., 2022), or participation in cell profiling adipocyte precursors (Burl et al., 2022). Of note, most trials studied macrophage dynamics in a context in which BAT was thermogenically activated through exposure of mice to low temperatures. Although some authors have suggested an increase in proinflammatory macrophages in the BAT of mouse models of obesity, most works have confirmed the role of macrophages in impairing the functionality of white adipose tissue (WAT) (Kriebs, 2021). Through the use of scRNAseq methods, however, it emerges that a subclass of macrophages, commonly annotated as lipid-associated macrophages (LAMs), play a protective role in limiting the metabolic degeneration observed in both genetic and dietary models of obesity (Jaitin et al., 2019; Liu et al., 2019). It has also been reported that LAMs develop a foamy phenotype and increased metabolic activity and lysosomal mass (Coats et al., 2017; Xu et al., 2013), such as some authors have shown an increased capacity of LAMs to secrete proteins by virtue of their senescent phenotype (Rabhi et al., 2022; Sawaki et al., 2023). Taken together, these data suggest that there is still no common consensus on the role of BAT macrophages in conditions of obesity.

In order to provide a detailed picture on monocyte/macrophage dynamics in BAT in obesity, here we have developed MACanalyzeR, a multilayered scRNAseq based computational tool describing the foamy-like features, polarization status and metabolic projection. In line with the data reported for white adipose tissue, in the BAT of obese mice we observed a significant increase in the cluster pertaining to LAM which showed a significant enrichment of genes belonging to mitochondria and lysosomes. The detailed analysis of phenotypic and metabolic parameters reports that BAT LAMs in diet-inducing obesity models develop peculiarities associated with the PPAR_γ_ signaling pathway. We then traced the dynamics of *Pparg*^HIGH^ LAMs in notexin-injured skeletal muscle, and observed a significant accumulation during the regenerative phases.

In conclusion, we can suggest MACanalyzeR as a versatile computational tool to dissect the macrophage responses from data obtained by scRNAseq. Our predictions highlight *Pparg*^HIGH^ LAMs as a metabolically active subclass potentially contributing to the regenerative process of BAT thus maintaining its thermogenic potential.

## RESULTS

### Lipid-associated macrophages accumulate in BAT of murine models of obesity

BAT serves as a crucial metabolic regulator, facilitating the clearance of circulating nutrients to meet its bioenergetic demands during non-shivering thermogenesis. BAT activity is significantly reduced in both mice and humans with type 2 diabetes. While many studies have discussed the homeostatic role of macrophages in BAT during physiological conditions like cold exposure, the behavior of macrophages in conditions deactivating BAT such as obesity remains largely unexplored.

To address this gap, we aimed to investigate the phenotypic response of brown adipose tissue (BAT) of mice with type 2 diabetes (db/db) and those fed a high-fat diet (HFD). We initially analyzed bulk RNA sequencing (RNAseq) data from the BAT of db/db and HFD mice, focusing on the expression levels of canonical brown or white fat genes. Notably, the BAT of db/db mice exhibited a reduced identity of brown adipocytes and a more white-like phenotype compared to HFD mice, which preserved their brown fat identity **(Fig. 1A)**. This transition from BAT to WAT gene expression in db/db mice may underlie the loss of thermogenic activity, potentially contributing to the observed metabolic dysfunction and obesity **(Fig. 1B-C)**.

**Figure 1.**
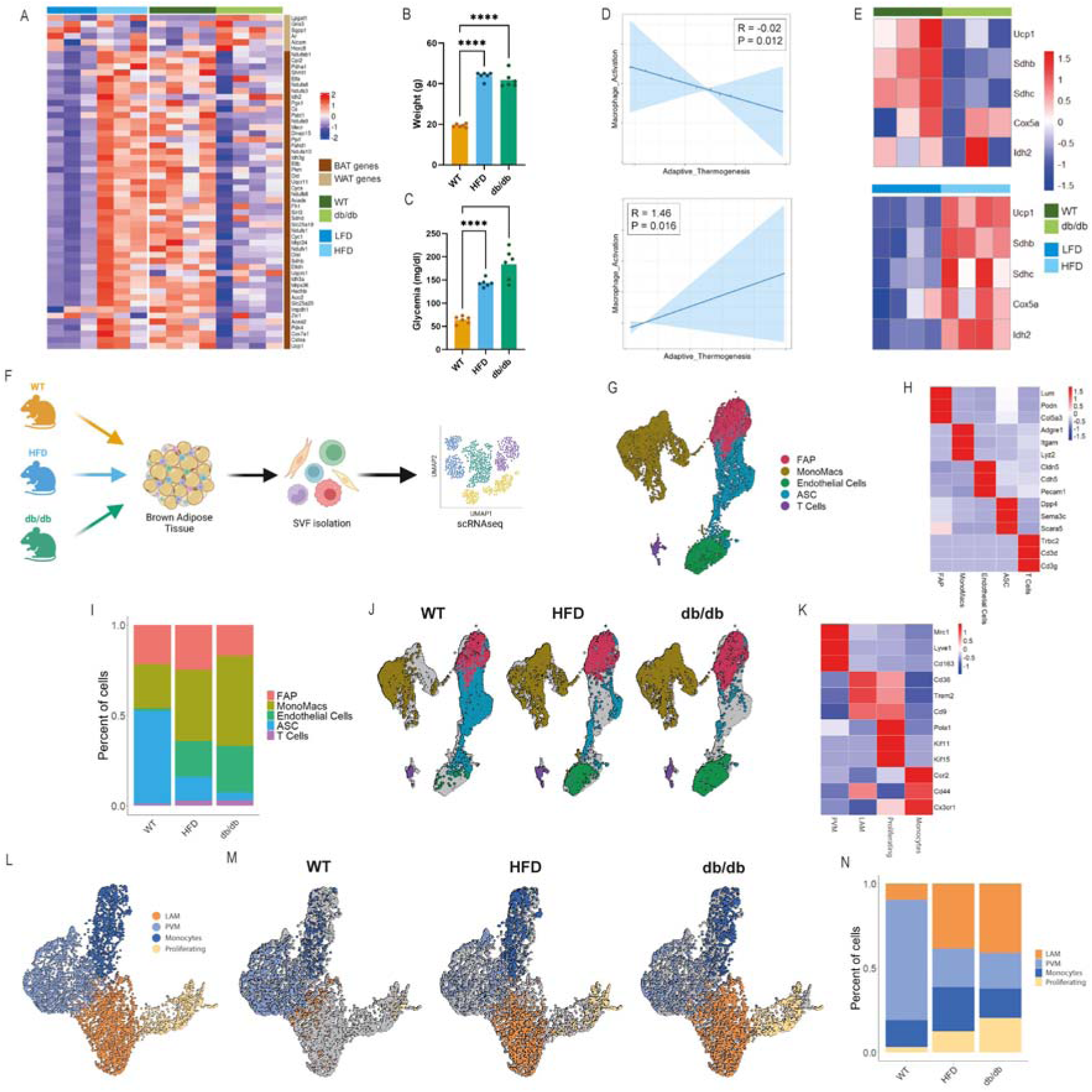
scRNA analysis reveals LAM accumulation in BAT of obese mouse models. (**A**) Heatmap of BAT and WAT markers in bulk RNAseq data of HFD and db/db BAT. (**B-C**) Barplot reporting body weight of db/db and HFD mice, data were reported as mean SD. ANOVA test **p<0.01. (**D**) Heatmap of thermogenic proteins in proteomics data of db/db (*upper panel*) HFD (*lower panel*) BAT. (**E**) Linear models fitted on scores computed on GO terms Macrophage Activation pathway and Adaptive Thermogenesis for BAT db/db data (*upper panel*) and BAT HFD data (*lower panel*). (**F**) Graphical cartoon of scRNAseq experimental design. (**G**) UMAP representing cell clusters identified by scRNAseq of SVFs (n=14715) isolated from BAT of wild type (wt), db/db and high fat diet (HFD)-fed mice. Clustering identified 5 major clusters: Fibro-Adipose Progenitors (FAP), Adipose Stromal Cells (ASC), Endothelial Cell, T cells and Monocytes/Macrophages (MonoMacs). (**H**) Heatmap of cell-marker genes expression in cell clusters. (**I**) Barplot representing the cell proportion across the 3 conditions. (**J**) UMAP of BAT SVF splitted by conditions. (**K**) Heatmap reporting marker-gene expression of MonoMacs subclusters. (**L**) UMAP of MonoMacs sub-clustering (n=5578) identified by scRNAseq analysis. Clustering identified 4 major clusters: Monocytes, Perivascular Macrophages (PVM), Lipid-Associated Macrophages (LAM) and Proliferating Macrophages. (**L**) UMAP of BAT MonoMacs splitted by condition. (**M**) Bar plot of MonoMacs subclusters abundance across the 3 conditions.

To strengthen the hypothesis that macrophages may play a role in the loss or maintenance of thermogenic function and brown adipocyte identity, we computed our pathway score for the GO pathways (Materials and Methods) related to thermogenic function and macrophages activation for each sample. Subsequently, we constructed two linear models based on these scores **(Fig. 1D)**, which led to the identification of an inverse relation between macrophage activity and thermogenic function in db/db mice. Interestingly, the HFD mice showed a positive correlation between these two scores, suggesting that under HFD LAMs sustain BAT activity. This result was also confirmed by proteomics data that highlight a decrease of thermogenesis in the BAT of db/db mice, whereas BAT from HFD mice showed induced thermogenesis **(Fig. 1E)**.

Next, we aimed to investigate the cell heterogeneity in BAT of db/db and HFD mice and gain a deeper insight into the dynamics and transcriptomic changes in immune cells, specifically in macrophages To achieve this, we analyzed single-cell RNA sequencing (scRNAseq) data collected from stromal vascular fraction (SVF) isolated from BAT in both db/db and HFD-induced obese mice (**Fig. 1F**). After rigorous quality filtering, the analysis led to the identification of a total of 14715 cells (∼4900 cells sampled per condition: WT, db/db and HFD). Based on differential gene expression and well-known literature markers, we identified 6 distinct clusters that were annotated as Fibro-Adipose Progenitors (FAPs), Adipose Stromal Cells (ASCs), Endothelial

Cells, T Cells and monocytes/macrophages (MonoMacs) **(Fig. 1G-H).** Of note, among all immune cells the MonoMacs cluster is noticeable the largest in cell number in BAT **(Fig. 1I)**. The population of MonoMacs massively increased in BAT in both db/db and HFD mice when compared with controls **(Fig. 1J)**. Importantly, a recent study highlights that scRNAseq experiments may exhibit biases towards specific cell types (Denisenko et al., 2020). Consequently, while we acknowledge the potential presence of technology-mediated biases in our data, the strong reproducibility observed across the two obese conditions and other studies (Jaitin *et al*., 2019) indicates that our compositional estimates remain largely unaffected by such technical factors.

In order to give more insight on MonoMacs dynamic in obese BAT, we sub-clustered and annotated these cells (n = 5578) using well-known biomarkers of AT macrophages from the literature **(Fig. 1K)**. This analysis led to identification of four distinct subclusters **(Fig. 1L)**, including one composed of monocytes expressing *Ccr2, Cd44, Cx3cr1*, one containing Lyve1^HIGH^ and Mrc1^HIGH^ macrophages denoted as perivascular-like macrophages (PVMs), and two with a very similar expression patterns, which can be attributed to the previously reported lipid-associated macrophages (LAMs). Among these two latter groups, one is characterized by genes involved in cellular proliferation (*Pola1, Kif11, Kif15*), leading us to designate these macrophages as proliferating macrophages (proliferating).

In line with the notable accumulation of LAMs in expanding WAT and BAT (Duerre and Galmozzi, 2022; Jaitin *et al*., 2019), a significant increase in LAMs was also observed in BAT of db/db and HFD mice **(Fig. 1M-N)**. These macrophages shifted from being almost absent in control mice to becoming the most abundant cluster in obese mice.

### MACanalyzeR identifies foamy-like macrophages in obese BAT

Considering the growing evidences highlighting the pivotal role in macrophages in maintaining tissue homeostasis, and the current gap in single-cell computational tools specifically designed to analyze both the transcriptomic and metabolic profiles of these cells, we introduce MACanalyzeR. MACanalyzeR is designed to characterize monocytes and macrophages in single-cell transcriptomics with 3 modules that address key aspects of these cells such as foamy-like features (FoamSpotteR), polarization status (MacPolarizeR) and metabolic phenotype (PathAnalyzeR). We applied MACanalyzeR to deconvolute the metabolic and phenotypic heterogeneity of brown adipose tissue macrophages under obesity conditions.

Previous studies have underscored the crucial role of foam macrophages (fMACs) in atherosclerotic plaques, modulating inflammation and releasing key factors that stimulate cell proliferation and differentiation, thereby facilitating the regeneration process (Fernandez and Giannarelli, 2022). However, a clear description of these foamy-like features in adipose tissue macrophages is lacking. Furthermore, while the development of this phenotype in macrophages within obese WAT has been documented, its occurrence in BAT remains unexplored. To thoroughly decipher the foamy-like features in BAT macrophages, we developed FoamSpotteR, a module based on a machine learning classifier that utilizes cellular transcriptional profiles at a single-cell level **(Fig. 2A)**. To train this model, we analyzed scRNAseq data collected from aortic CD45^+^ (*Ptprc*) cells isolated from healthy mice and atherosclerotic mice from which we extracted macrophages and annotated them as fMAC^-^ and fMAC^+^ based on specific markers found in literature (Cochain et al., 2018; Kim et al., 2018) **(Fig. 2B, Fig.S1A-D)**. For model training, we utilized 75% of the total cells, while the remaining 25% were used as a test set. The model demonstrated high efficiency and very high accuracy in discriminating fMAC^+^ in single-cell data.

**Figure 2.**
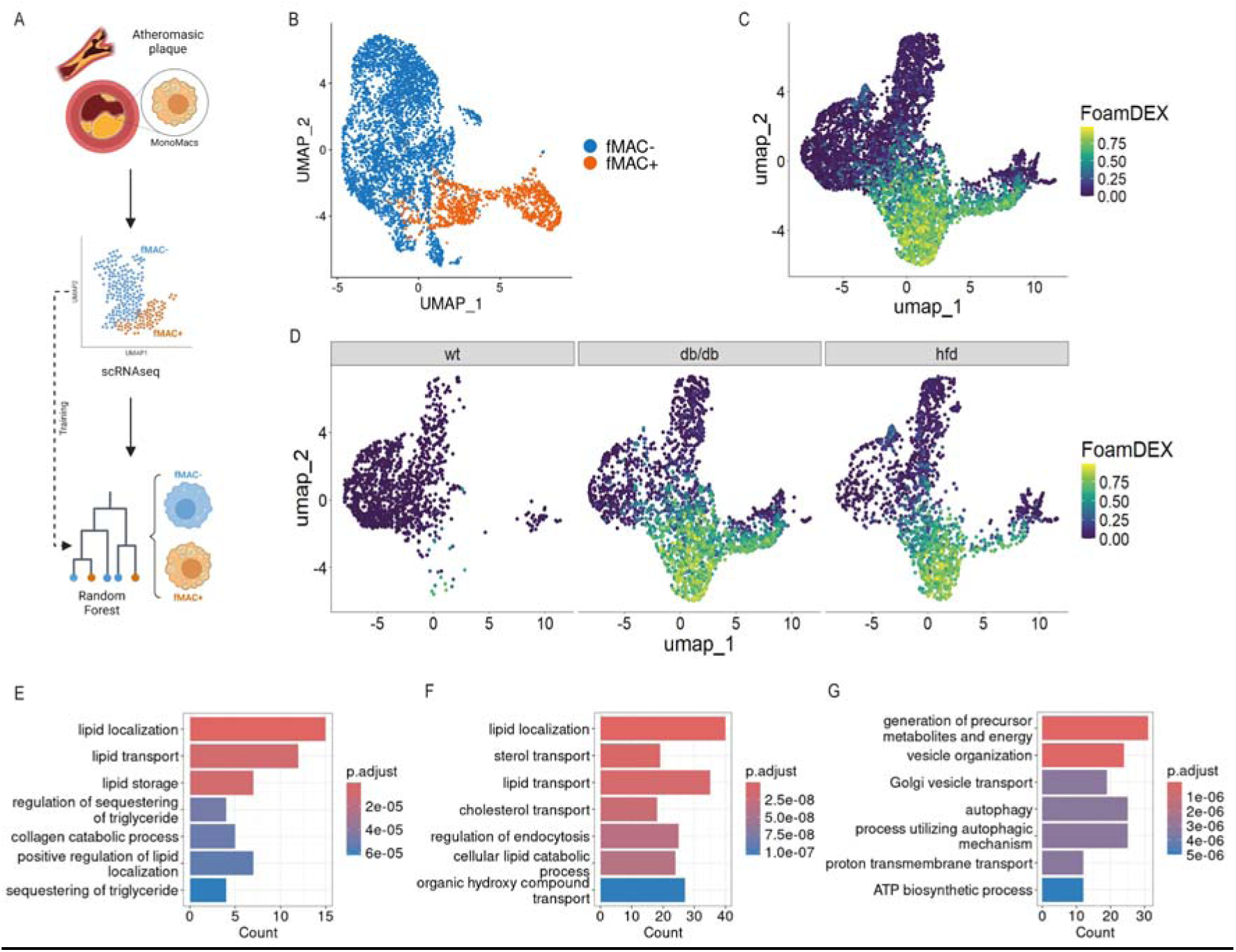
FoamSpotteR identifies foamy-like macrophages in obese BAT. (**A**) Graphical representation of the FoamSpotteR module workflow. (**B**) UMAP of macrophages isolated from atheromatous plaque (GSE97310 and GSE116240). Clustering identified two major clusters: foam macs (fMAC^+^) and non-foam macs (fMAC^-^). (**C-D**) UMAP of BAT macrophages colored by FoamSpotteR result (C) and splitted by sample (D). Values above 0.5 of FoamDEX identify foamy-like macrophages. (**E-G**) Gene Set Enrichment for GO Biological Process of shared upregulated genes in bulk RNAseq of foam macrophages and FoamSpotteR predicted fMAC^+^ (E), uniquely upregulated in foam macrophages (F) and uniquely upregulated in fMAC^+^ predicted cells (G) (pval < 0.05; log2FC>1).

Next, we utilized the trained FoamSpotteR model to analyze macrophages in BAT and observed a notable presence of fMACs in both db/db and HFD mice **(Fig. 2C-D)**. To determine the emergence of fMACs within a specific macrophage subcluster, we calculated the proportion of fMACs in each subpopulation. As expected, we observed that the occurrence of fMACs predominantly corresponds to LAMs and proliferating macrophages, constituting 57% and 27% respectively.

To validate the predictions generated by FoamSpotteR and clarify the differences between fMACs and LAMs, we compared our results with those from a publicly available dataset from bulk RNA seq (GSE116239), which includes foamy and non-foamy macrophages isolated from atherosclerotic aortas via lipid probe-based flow cytometry sorting (Kim *et al*., 2018). We conducted a comparative analysis by examining the Differentially Expressed Genes (DEGs) of fMACs from the bulk RNAseq and those identified through our FoamSpotteR module in the previously annotated LAM subcluster (fMAC^+^ LAMs vs. fMAC^-^ LAMs). Our investigation revealed 593 upregulated genes in fMACs from bulk RNAseq and 193 in fMAC^+^ LAMs from our scRNAseq, with 73 genes shared between the two datasets.

Notably, the commonly upregulated pathways are associated with lipid metabolism and localization **(Fig. 2E)**. In contrast, pathways uniquely upregulated in fMACs from atherosclerotic plaque predominantly involve cholesterol metabolism **(Fig. 2F)**. Interestingly, the pathways uniquely upregulated in fMACs in BAT are linked to lysosomes and mitochondria **(Fig. 2G)**. This suggests an overlap between the two cell types, yet indicates a metabolic distinction that renders these cell types metabolically distinct despite their phenotypic similarities.

### BAT LAMs show a different polarization status between db/db and HFD mice

Traditionally, macrophages are classified into three different categories according to their activation status: inactivated/quiescent macrophages (M0), classically activated (M1, pro-inflammatory) and non-classically activated (M2, anti-inflammatory) (Yunna et al., 2020). While AT macrophages (ATMs) exhibiting an M1-like phenotype are well-established contributors to impaired insulin sensitivity in obese mice and humans, some studies have reported that ATMs isolated from the AT of obese individuals display a less inflammatory profile compared to classically activated M1 macrophages, (Coats *et al*., 2017; Rosina *et al*., 2022).

Based on this evidence, we hypothesize that the observed difference in polarization phenotypes might be due to inherent variability within the ATM population. To investigate this hypothesis, we developed MacPolarizeR, a machine learning-based module that classifies macrophages according to their activation state. MacPolarizeR operates by clustering cells according to the expression of genes associated with M1 and M2 polarization, selected from a public dataset (GSE129253) including bulk RNAseq data of M1 and M2 macrophages polarized in vitro. Following this training process, MacPolarizeR can identify three distinct macrophage populations: Inflammatory (*M1-like*), Transitional (*M0-like*), and Healing (*M2-like*).

In addition to this classification, MacPolarizeR calculates an M1 and M2 polarization score for each cell based on the expression of the previously selected genes. This score system facilitates a more comprehensive and nuanced characterization of the macrophage polarization state. The calculated scores can be visualized on a two-dimensional plot, with M1 score on the *X-axis* and M2 score on the *Y-axis* **(Fig. 3A)**. This graphical representation provides a clearer interpretation of macrophage behavior, indicating whether subpopulations lean towards a healing phenotype (upper left), a transitional state (upper right and lower left), or an inflammatory state (lower right).

**Figure 3.**
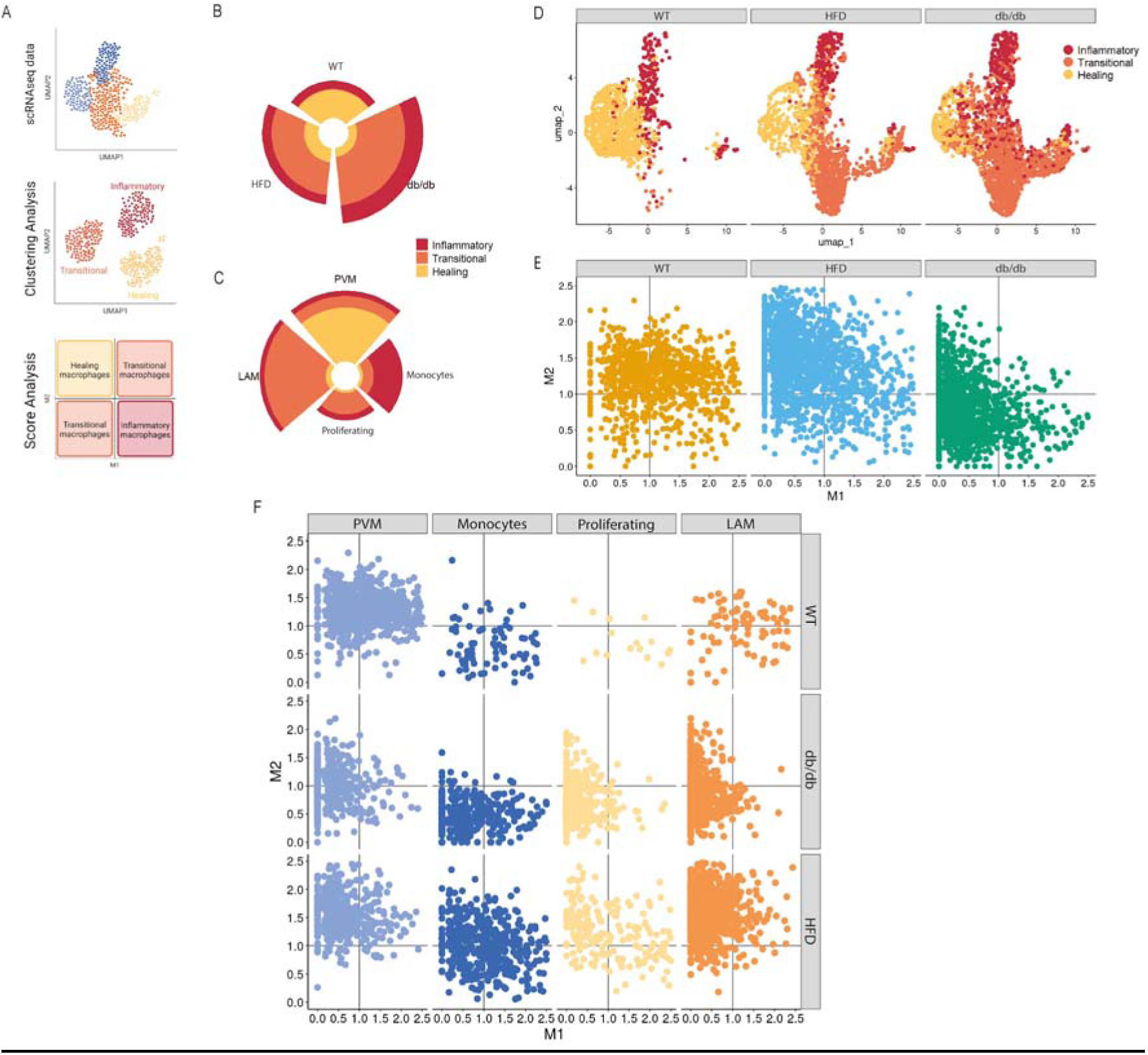
MacPolarizeR shows a less inflammatory phenotype of LAMs in obese BAT. (**A**) Graphical cartoon of MacPolarizeR workflow. (**B-C**) Circular bar plot representing MacPolarizeR’s classification across conditions (B) and macrophage subcluster (C). (**D**) UMAP of BAT macrophages colored by MacPolarizeR result and splitted by conditions. (**E-F**) 2D-plot of MacPolarizeR M1 and M2 score across conditions (E) and across conditions and macrophage subclusters (F).

To validate the accuracy of our polarization classification, we applied MacPolarizeR to a scRNA sequencing dataset derived from *in vitro* polarized macrophages (GSE117176). Specifically, bone marrow derived macrophages (M0 phenotype) were treated with IFN-_γ_ and LPS to induce an M1 phenotype, and IL-4 plus IL-13 to induce an M2 phenotype. The analysis of this dataset gave rise to the identification of 3 distinct clusters based on the forced polarization status that we used as input for MacPolarizeR **(Supplementary** Fig.2A). Although the identification of inflammatory macrophages was very precise, there was an overlap between transitional and healing macrophages related to the similarity between these two subclasses, as the original authors pointed out (Li et al., 2019) **(Supplementary** Fig.2B). Noteworthy, the polarization score analysis highlights a less pronounced overlap between M0 and M2, confirming that the prediction made by MacPolarizeR perfectly aligns with the polarization observed *in vitro* **(Supplementary** Fig.2C).

For further validation, we applied MacPolarizeR to analyze a scRNAseq dataset derived from the WAT of mice at one day and one-month post-sepsis induction (PRJNA626597), a representative model system for studying *in-vivo* inflammation **(Supplementary** Fig.2D). This dataset represents a perfect fit for our module, as it includes two different inflammatory status: acute (one day post sepsis) and chronic (one month post sepsis). After the scRNAseq analysis, the MonoMacs cluster was extracted from the dataset and used as input for MacPolarizeR **(Supplementary** Fig.2E). As expected, after one day of sepsis the macrophage component shifted towards an inflammatory phenotype (M1). In contrast, after one month of sepsis, the macrophages acquired a completely different phenotype, predominantly residing in the M2 quadrant of the MacPolarizeR plot, indicative of healing macrophages. In contrast, macrophages in control WAT exhibited an intermediate localization on the graph, suggesting an inflammation-suppressed phenotype **(Supplementary** Fig.2F**-G).** The use of MacPolarizeR not only provided insights into the activation states of macrophages but also demonstrated its effectiveness as a tool for offering a more granular and clearer depiction of specific phenomena. By using MacPolarizeR, we were able to trace a sequential activation of macrophages from early to late time points following a specific stimulation. This progression began with a predominantly-inflammatory contribution during the acute phase of the pathology, which then transitioned to a healing state.

Considering the high accuracy achieved, we applied the MacPolarizeR module to the single-cell profile of BAT macrophages to dissect the differences between overfeeding (db/db) and HFD phenotypes. This approach aimed to enhance our understanding of how the tissue either maintains homeostasis or undergoes degeneration. Consistent with the increased macrophage component in BAT of the two obese models, our clustering analysis revealed a significant rise in transitional macrophages, a phenomenon that was almost absent in BAT of control mice. Specifically, we observed that the increase in transitional macrophage components coincided with the accumulation of LAMs in both db/db and HFD mice. In contrast, monocytes exhibited inflammatory behavior, while PVMs primarily displayed healing properties **(Fig. 3B-D)**.

To refine polarization characteristics, we conducted a scoring analysis to assess M1 or M2 polarization. Interestingly, LAMs of HFD mice revealed the acquisition of a healing phenotype in contrast to the LAM of db/db mice, which exhibited a transitional phenotype **(Fig. 3E-F)**. The healing polarization of macrophages in HFD mice suggests a potentially different response of BAT compared to that from db/db mice.

### Foam LAMs showed an enhanced mitochondrial and lysosomal metabolism

Next, we asked if fMACs in BAT of db/db and HFD mice undergo to metabolic reprogramming that is functional to their function. To explore the metabolic phenotype of BAT macrophages, we developed a novel scoring system that compares the expression levels of pathway-related genes under various conditions. This system incorporates cell-specific gene expression, allowing for a more precise analysis of metabolic changes. This approach departs from traditional techniques like over-representation analysis, which neglects cellular expression data, and Gene Set Enrichment Analysis (GSEA), which can be influenced by genes with low or no expression. Our method calculates a pathway activity score by averaging the weighted gene expression of all genes within a pathway across a specific cell type. This score, applicable at both the condition and cell cluster level, provides a clear indication of pathway upregulation (score > 1) or downregulation (score < 1).

We started by analyzing KEGG pathways associated with metabolism, specifically focusing on those with roles or components unique to macrophages. By comparing the conditions, we observed that BAT macrophages in db/db and HFD mice displayed upregulated pathways related to the pentose phosphate pathway, glycolysis, TCA cycle and Mitochondria (**Fig. 4A**). Remarkably, these results highlighted a higher metabolic activation of LAMs than other MonoMacs (**Fig. 4B**). However, a key difference emerged between the two models: db/db macrophages exhibited a prominent shift towards oxidative phosphorylation (OXPHOS), while HFD macrophages primarily displayed increased lipid metabolism and lysosome upregulation. To validate these findings, we generated heatmaps for each condition depicting genes belonging to these three pathways, which confirmed the results obtained from PathAnalyzeR (**Fig. 4C-D**).

**Figure 4.**
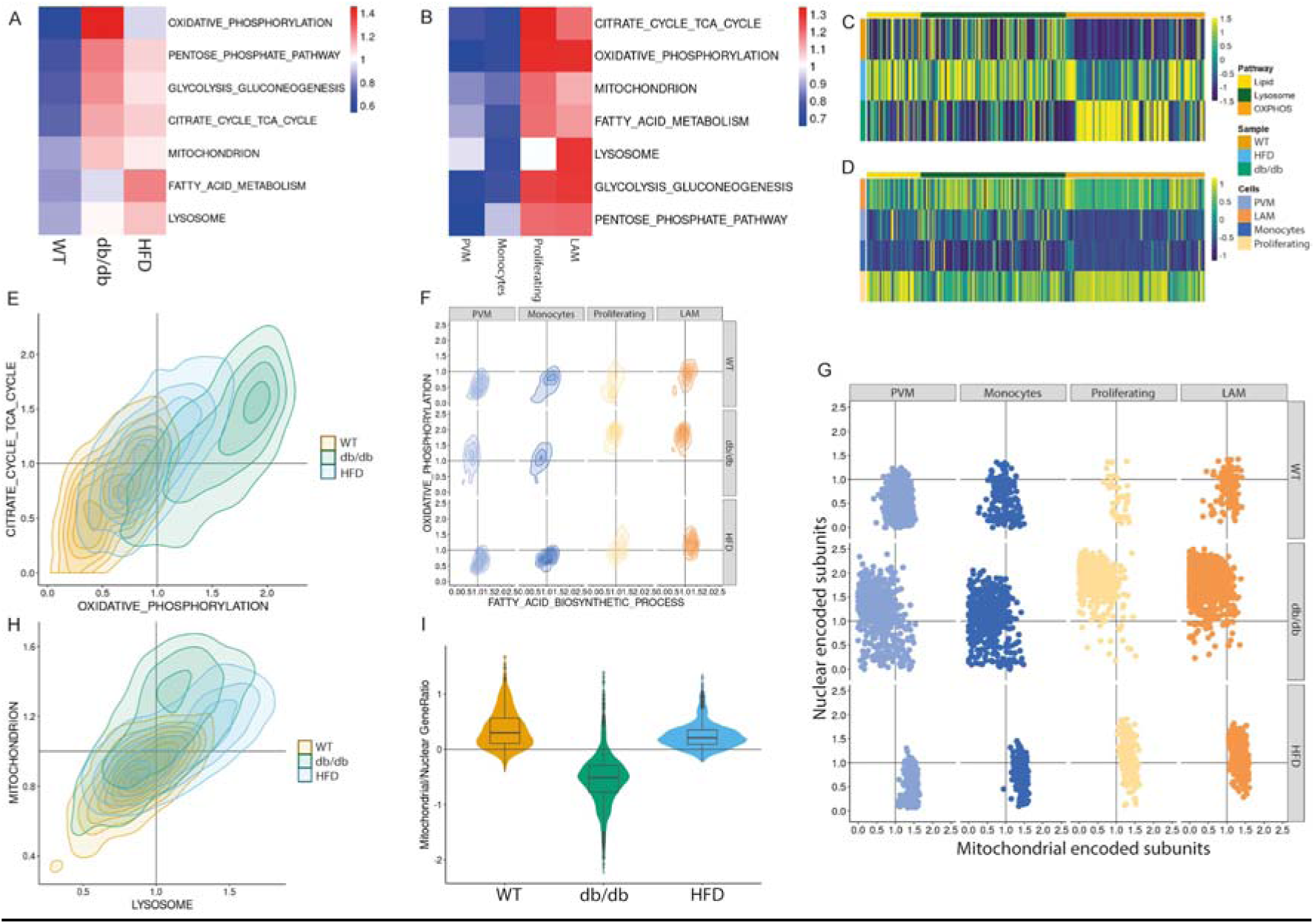
PathAnalyzeR describes metabolic differences in BAT LAMs of db/db and HFD mice. (**A-B**) Heatmaps of PathAnalyzeR analysis using KEGG metabolic pathways related to roles or components unique to macrophages. Plotted by condition (A) and by macrophage subcluster (B). (**C-D**) Heatmap representation of gene expression of Lipid Metabolism, OXPHOS and Lysosome pathways. Plotted by condition (C) and by macrophage subclusters (D). (**E-F-G-H**) 2D plot of PathAnalyzeR single-cell analysis. (**I**) ViolinPlot of MitoNuclear imbalance in BAT macrophages

One of PathAnalyzeR’s additional features is its ability to calculate pathway scores at the single-cell level. This enables a more refined metabolic characterization of each cell cluster, providing a deeper understanding of enriched pathways within the expression profile. This functionality allows for the simultaneous visualization of two distinct pathways on a two-dimensional density plot, facilitating comparisons between conditions or cell types.

To observe whether the up-regulations were consistent, we calculated single-cell scores for pathways related to OXPHOS and TCA cycle **(Fig. 4E)**. The density plot also reveals that db/db macrophages have high oxidative metabolism, whereas those from db/db and HFD undergo a shift towards enhanced TCA cycle activity compared to the control group. To clarify the type of lipid metabolism alteration operating in the two conditions, we computed pathway enrichment scores for specific Gene-Ontology (GO) terms related to lipid metabolism (i.e., catabolic vs biosynthetic). Notably, while there were no significant differences in fatty acid catabolism between the two conditions, our analysis revealed an upregulation of lipid biosynthetic processes in HFD macrophages **(Fig. 4F)**.

To investigate whether the different regulation of these pathways was attributable to a specific cell type, we plotted these two pathways separated by both condition and cell type **(Fig. 4G)**. Interestingly, the up-regulation in both cases is primarily attributed to LAMs, although they exhibit different patterns, with db/db LAMs distinctly oriented towards OXPHOS pathways, while HFD LAMs more engaged in fatty acid biosynthesis.

The differential metabolic response of identical cell types could be due to the varying obesity-inducing mechanisms between db/db overfeeding and HFD. This divergence in metabolic phenotypes of macrophages may correlate with the maintenance of brown adipocyte identity that we observe in BAT HFD and that is totally lacking in BAT db/db.

To further investigate whether major MonoMacs cellular components are also differentially regulated, we calculated single-cell scores for mitochondria and lysosomes and plotted them in a 2D scatter plot (**Fig. 4H**). It can be observed that in both db/db and HFD mice, there is a significant increase in either the lysosomal or mitochondrial components compared to controls. To understand the origin of the increased mitochondrial component, we also calculated the MitoNuclear imbalance score, which is the ratio of the score related to the expression of mitochondrial genes to the expression of nuclear-encoded mitochondrial genes (**Fig. 4I**). Interestingly, it is evident that the increase in mitochondrial components in db/db and HFD mice is due to different mechanisms. In the case of db/db mice, this increase is due to a shift in the nuclear-encoded mitochondrial components, while in HFD mice to an increase in both mitochondrial and nuclear components, as there is a small shift compared to controls.

### PPAR**_γ_** as a potential driver of regenerative macrophages

To further explore the transcriptional trajectories that committed macrophages towards these two types of activation (healing in HFD and transitional in db/db), we performed velocity analysis on BAT MonoMacs from both db/db and HFD models. The resulting vectors were projected onto the respective UMAP plots **(Fig. 5A-B)**. Consistent with our expectations, the velocity projections suggest that LAMs arise from the differentiation of monocytes and PVMs. The key difference between the two models is observed in the monocyte velocity. In particular, the db/db model shows a lack of directional information for monocyte vectors, which is present in the HFD model. Interestingly, the directionality exiting within PVMs remains consistent across both BAT conditions.

**Figure 5.**
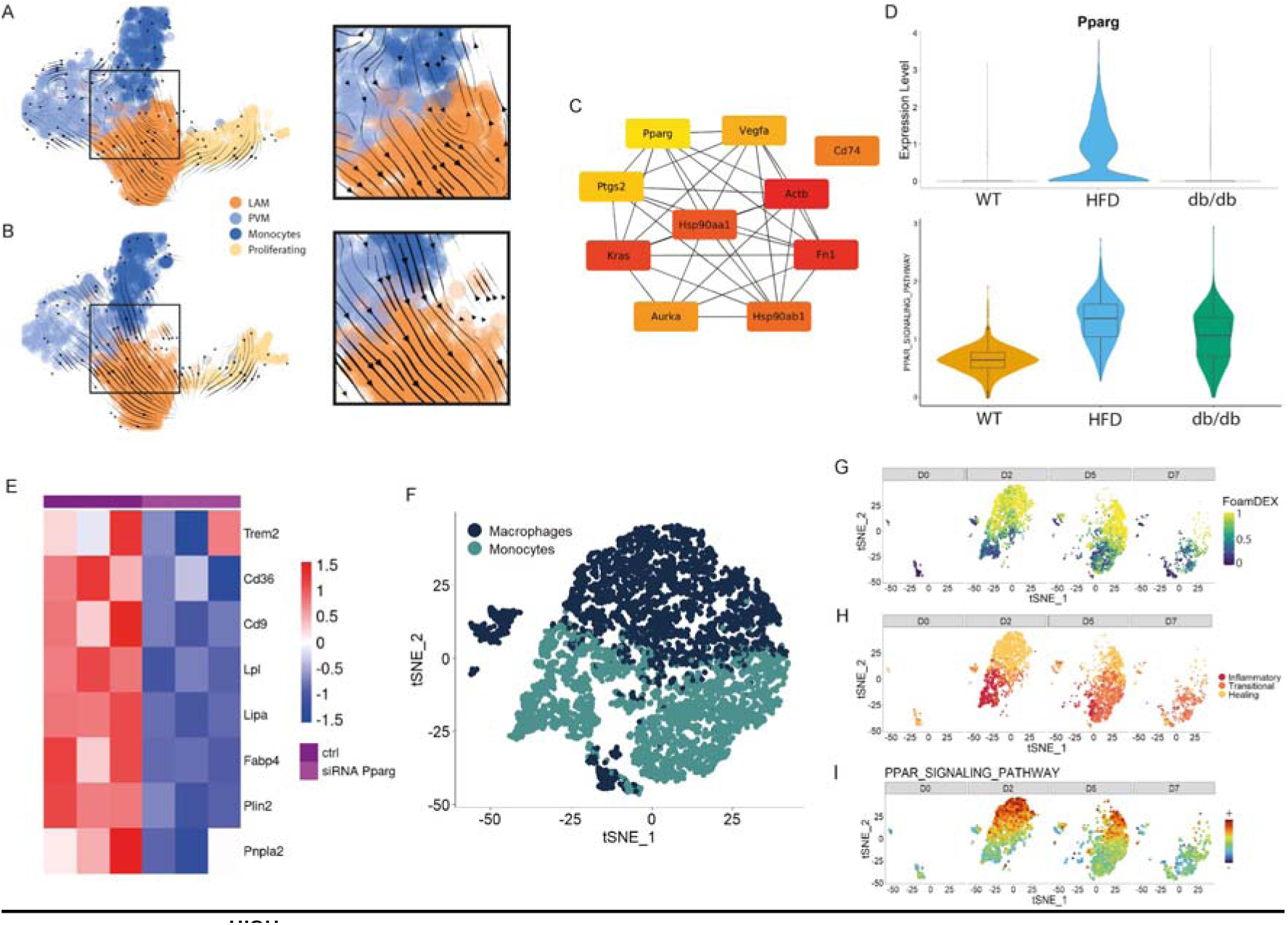
*Pparg*^HIGH^ LAMs contribute to tissue regeneration. (**A-B**) scRNAseq velocities projected onto the UMAP in db/db MonoMacs (A) and HFD MonoMacs (B). (**C**) HubGenes of the PPI network of HFD velocity genes. (**D**) Expression of *Pparg* (*upper panel*) and PPARγ signaling pathway (*lower panel*) splitted by conditions. (**E**) Heatmap of LAM signature gene expression in IL4-treated macrophages, ctrl and PPARγ downregulated. (**F**) UMAP representing MonoMacs subclusters identified by scRNAseq of Notexin-treated skeletal muscle (n=7572) isolated after 0, 2, 5 and 7 days after injury (GSE143437). (**G-H-I**) Result of MACanalyzeR analysis on skeletal muscle’ MonoMacs. Result of FoamSpotteR (G), MacPolarizeR (H) and PathAnalyzeR score of PPAR signaling (I) projected onto the UMAP.

The observed differences in LAMs between the two models suggest distinct differentiation patterns. Notably, the pronounced healing and lipidic biosynthetic behavior of LAMs in HFD conditions could be attributed to a unique molecular differentiation mechanism in monocytes that is entirely absent in the db/db model. This difference suggests precise and distinct molecular processes underlying the observed behaviors.

Velocity analysis enables us to identify critical genes for differentiation by ranking their contribution to velocity calculations. By focusing on genes relevant to differentiation, we can link the observed behavior to a potential molecular mechanism. Since the differences between db/db and HFD models were observed in the monocyte cluster, we restricted the gene ranking analysis to this subpopulation. This analysis identified 481 genes in db/db and 345 in HFD, with 90 genes shared between the two models.

To elucidate the molecular basis of the distinct polarizations and metabolic phenotypes, we identified the top 10 hub genes among the 345 unique genes differentially expressed in the HFD group **(Fig. 5C)**. Notably, this set included *Pparg*, a well-known gene regulating the anti-inflammatory response, fatty acid biosynthesis, as well as mitochondrial OXPHOS (Charo, 2007; Zhang and Chawla, 2004). Interestingly, *Pparg* expression was significantly elevated only in HFD mice **(Fig. 5D)**. The hypothesis that the different behaviors may be connected to the PPAR_γ_ signaling pathway is further confirmed by the PathAnalyzeR module, as an upregulation of *Pparg* was detected in HFD.

To further elucidate the contribution of macrophage PPAR_γ_ to the development of LAM signatures and their characteristic foamy phenotype we analyzed publicly available RNAseq data. This data were compared with wild-type (ctrl) macrophages and macrophages with PPAR_γ_ downregulation, both stimulated with IL-4 **(Fig. 5E)**. The data revealed a robust enrichment of genes associated with LAM features upon IL-4 treatment. Conversely, PPARγ deficiency resulted in a marked downregulation of these LAM signature genes. These findings provide strong evidence for the involvement of PPARγ in regulating the macrophage response to stimuli like IL-4, potentially influencing their differentiation towards LAM.

The collected results led us to postulate that LAMs with high Pparg expression (*Pparg*^HIGH^) take part in tissue regeneration. To demonstrate that we analyzed scRNAseq data of Notexin-treated skeletal muscle (GSE143437), a well-established regenerative model (Walter et al., 2023). We considered injured muscles at 0, 2, 5 and 7 days Notexin post-injection. Our analysis focused on the MonoMacs component. **(Fig. 5F)**.

Our analysis showed that there is a significant increase in MonoMacs at specific time points, i.e., at 2 and 5 days after injury. Analysis by MACanalyzeR revealed that the macrophages present at the peak of the injury exhibit a foamy-like phenotype similar to that occurring in BAT **(Fig. 5G)**. In addition, the MacPolarizeR module identified significant inflammatory polarization of monocytes 2 days post-injury. In contrast, the macrophage component displayed a healing behavior in the subsequent days, underscoring their participation in tissue repair **(Fig. 5H)**. Following injury recovery, we observed a general decrease in both foam and healing macrophages, with the system nearly returning to its baseline steady-state.

Regarding the *Pparg* expression, we also found evidence of its involvement in muscle regeneration. Notably, *Pparg* expression peaked at 2 days after Notexin injection and decreased after the regeneration process **(Fig. 5I)**. Homeostasis was restored in 7 days.

## Discussion

Macrophages are cells with high adaptive plasticity taking part in the homeostatic system of the tissue. BAT dynamically adapts its metabolism to sustain thermogenesis; however, although mounting evidence identifies macrophages as playing a key role in this process, few works have developed an in-depth analysis of their molecular and metabolic profile. Furthermore, most of the works focused on the macrophage responses of BAT of mice exposed to cold. Differently, few studies have analyzed macrophage dynamics in BAT of obese mice. Of note, contrasting results revealed that BAT showed activated metabolism following a HFD (also called diet-induced thermogenesis) and lower activity in type 2 diabetes. These contrasting findings led us to ask if monocyte/macrophage differences in BAT of db/db and HFD could occur. In order to solve this question, we have developed MACanalyzeR, a 3-in-1 tool based on the analysis of data from scRNAseq. Using a machine learning approach, MACanalyzeR returns information relating to the degree of foamy-like characteristics, polarization status and metabolic projections of macrophages resident in obese BAT. Consistent with the observations from WAT depots (Jaitin *et al*., 2019), BAT accumulates LAMs with a foamy-like phenotype. LAMs with foaming features showed a significant enrichment of genes pertaining to mitochondrial and lysosomal metabolism. Remarkably, the strong induction of the mitochondrial and lysosomal components was corroborated by development of pathway score at single-cell level. The high expression level of lysosomal and mitochondrial genes observed in LAMs of metabolically stressed BAT, suggests a potential adaptive metabolic process in macrophages that aligns with the clearance of extracellular debris (Coats *et al*., 2017), which could cause an accumulation of extracellular debris that led to the loss of brown adipocytes activity (Rosina *et al*., 2022). It is also emerging that LAMs could participate in BAT regeneration through direct communication with proliferating adipocyte precursors (Burl *et al*., 2022). Indeed, the increased mitochondrial TCA-related metabolites (e.g. succinate) or cholesterol esters could refill mitochondrial metabolism of proliferating cells (Mills et al., 2016). It is also interesting to suppose that LAMs could also produce heat through energy dissipating cycles (He et al., 2020). Consistent with this, lysosomal V-ATPase complex, which transports protons (H^+^) uses the ATP derived from the activity of mitochondrial lipid catabolism (Demers-Lamarche et al., 2016), and the hydrolysis of ATP by V-ATPase is an exothermic process (de Meis et al., 1997). Although the amount of heat generated by individual ATP molecules may be limited, the significant increase in mitochondrial and lysosomal detected in LAMs, accompanied by the significant increase in LAMs, could represent an integrated thermogenic system between uncoupled respiration of brown adipocytes and lysosomal ATPase of macrophages. In line with the protective role of LAMs, several authors reported that following HFD, depletion TREM2^+^ LAM exacerbates WAT dysfunction and metabolic abnormalities (Jaitin *et al*., 2019). Remarkably, TREM2^+^ LAMs exert a protective role in other metabolically-stressed tissues such as the heart (Piollet et al., 2024; Zhang et al., 2023).

However, MACanalyzeR highlighted metabolic divergences between BAT LAMs of db/db and HFD mice. The lower ratio of mtDNA-/nDNA-encoded genes suggests the occurrence of a mitonuclear imbalance in BAT LAMs of db/db mice. In particular, the higher expression level of nuclear-encoded genes, suggest a significant defect of mtDNA to sustain mitochondria functions. Accordingly, macrophages with lower mtDNA-encoded genes such as TFAM KO macrophages, showed a defective clearance of cellular debris (Gao et al., 2022). As a second point, BAT LAMs from HFD mice show a significant induction of genes pertaining to the PPARγ signaling pathway. Interestingly, several authors reported a regenerative potential of *Pparg*^HIGH^ macrophages (Varga et al., 2016), which is consistent with our evidence reporting a significant dynamic of LAM *Pparg*^HIGH^ in skeletal muscle regeneration. Although the role of LAM *Pparg*^HIGH^ in adipose tissue physiology is still unclear, disruption of PPARγ signaling pathway in myeloid cells impairs alternative macrophage activation, thereby predisposing these animals to development of diet-induced obesity, insulin resistance, and glucose intolerance (Odegaard et al., 2007). Keeping in mind the difference observed in BAT LAMs from db/db and HFD mice, we suppose that LAM *Pparg*^HIGH^ could maintain BAT cell identity promoting differentiation of adipocyte precursors to brown adipocytes. Differently, during type 2 diabetes BAT LAMs promote BAT-to-WAT transition forcing the differentiation of adipocytes precursors to white adipocytes.

In conclusion, we provided MACanalyzeR as a multilayered scRNAseq-based computational tool to deeply characterize macrophages. This is a versatile tool that can virtually predict macrophage dynamics, giving insight in development of foamy features, polarization/activation waves and metabolic changes, virtually in every pathophysiological condition and anatomic district. Our analyses on BAT macrophages led to identification of LAMs *Pparg*^HIGH^ as a subcluster of LAMs, which specifically could maintain BAT cell identity in the condition of diet-induced obesity.

---

**Supplemental Figure 1.**
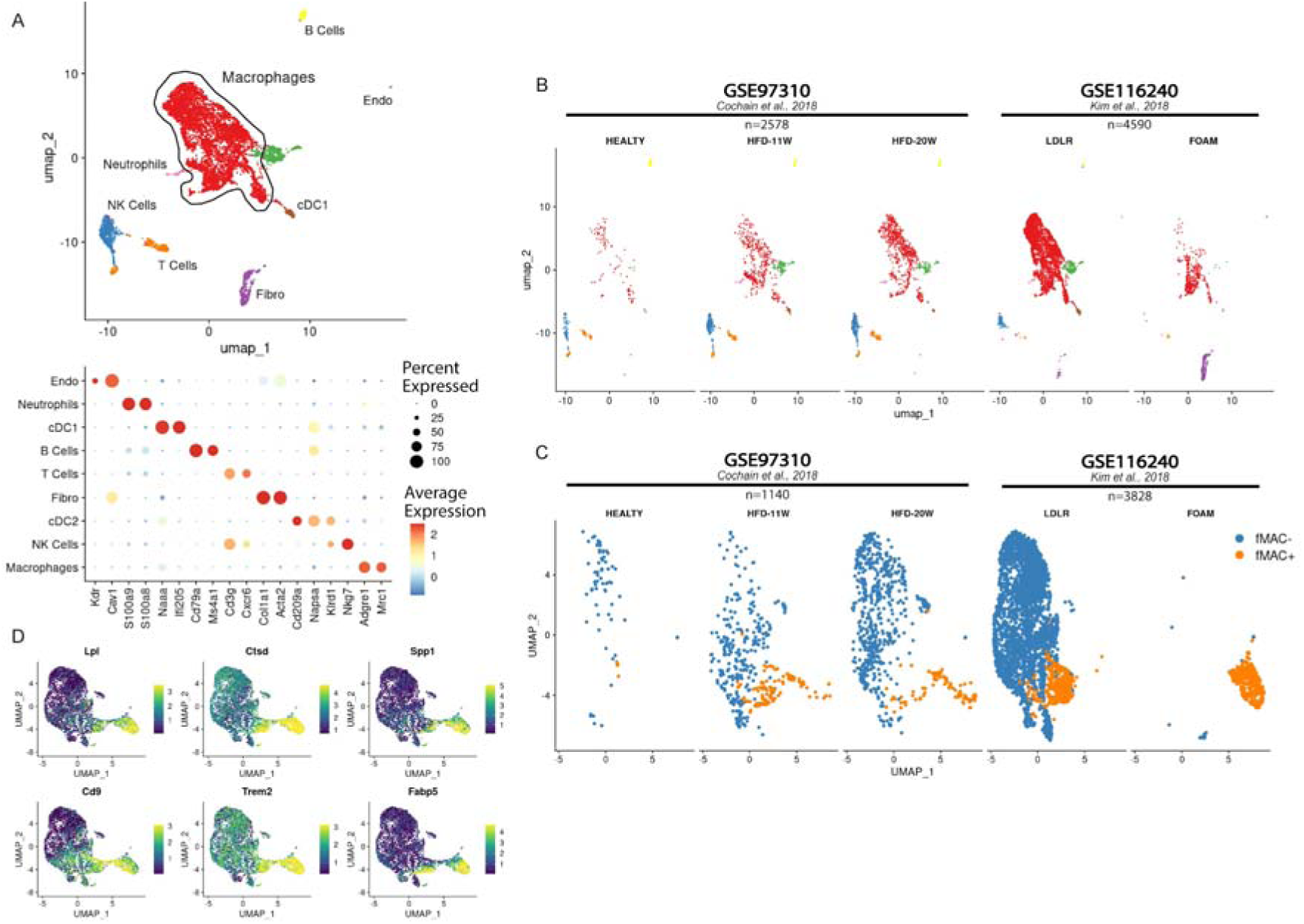
FoamSpotteR training set. (**A**) UMAP representing cell clusters identified by scRNAseq of CD45+ cells (n=7166) isolated from atherosclerotic aorta. Clustering identified 8 major clusters: Macrophages, Dendritic Cells (cDC1, cDC2), T Cells, B Cells, NK, Fibroblasts and Endothelial Cells (*upper panel*) and dotplot of cell markers expression (*lower panel*). (**B-C**) UMAP representation of each dataset further split according to conditions for total dataset (B) and macrophages subcluster (C). (**D**) Expression of transcript of the foamy cell signature projected onto the UMAP plot.

**Supplemental Figure 2.**
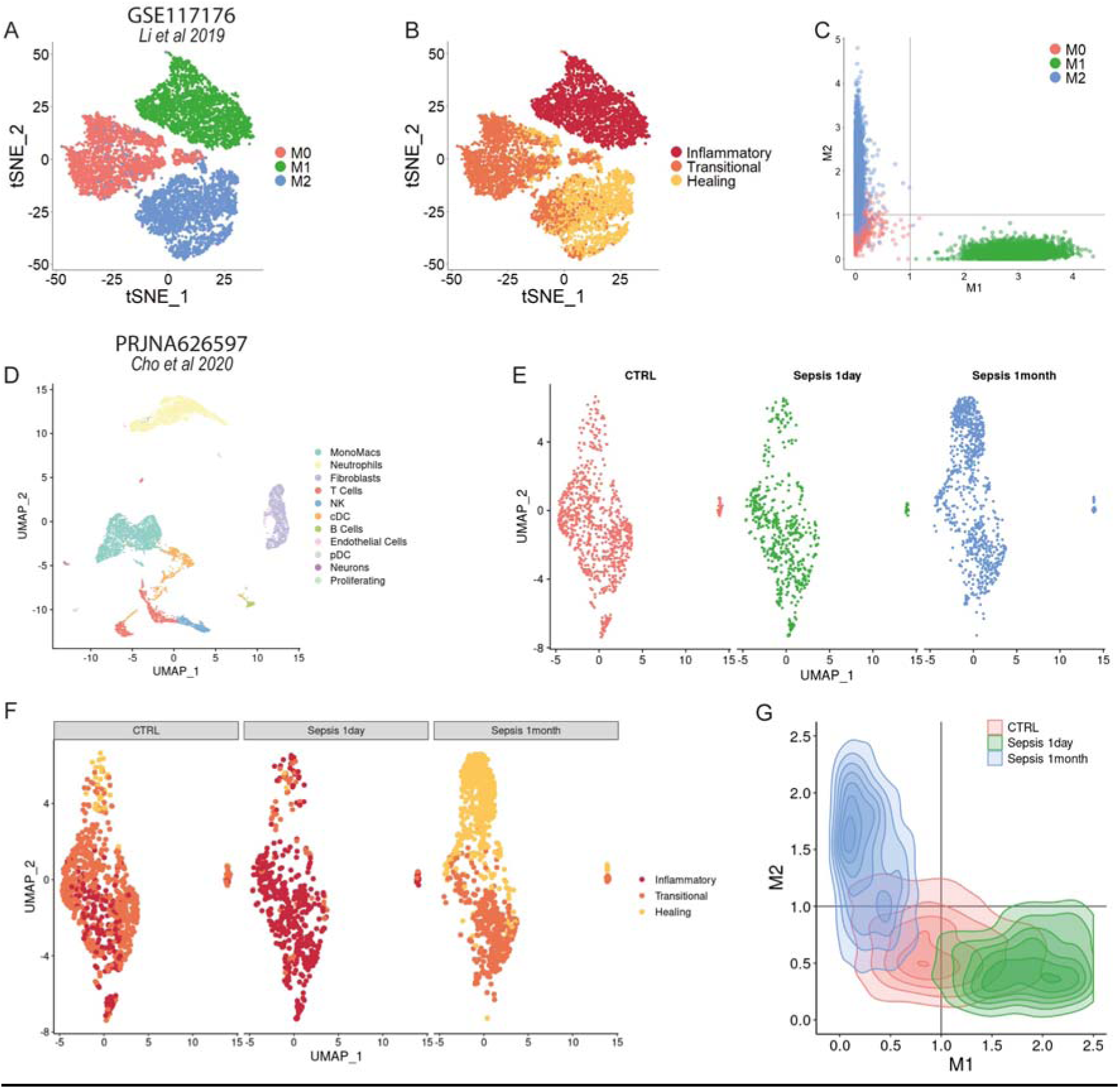
Validation strategy for MacPolarizeR module. (**A-B-C**) MacPolarizeR validation on scRNAseq data of in-vitro polarized macrophages. t-SNE of in-vitro polarized macrophages (A); the three clusters identified correspond to the induced polarization. t-SNE coloured by MacPolarizeR clustering prediction. (B). 2D plot of MacPolarizeR polarization scores (C). (**D-E**) UMAP representing cell clusters identified by scRNAseq of SVF (n=6900) of adipose tissue isolated from mice after 1 day of sepsis and 1 month of sepsis (D). Sub-clustering of monocytes and macrophages (E). (**F-G**) MacPolarizeR validation on scRNAseq data of septic adipose tissue. UMAP of MonoMacs clustered in septic adipose tissue, splitted by condition and coloured by MacPolarizeR clustering prediction (F). 2D plot of MacPolarizeR polarization scores (G).

## Limitation of the Study

Although MACAnalyzeR revealed interesting predictions data on the different responses of BAT macrophages, the study was lacking in some validation steps. In particular, the difference observed between BAT LAMs of db/db and HFD could be associated with treatment time. Although complex, it would in fact be desirable to do a time-course to better understand the dynamics of BAT LAMs during different phases of obesity. Furthermore, there is a lack of *in vitro* experiments to understand whether the metabolism of LAMs is higher if compared to non-LAMs. Finally, the regenerative role of *Pparg*^HIGH^ LAMs has not been tested in BAT.

## Acknowledgment

This work was mainly supported by the European Union - Next Generation EU, call “PRIN 2022 (Prot. 2022Z9HYJH)”, “PRIN 2022 PNRR (Prot. P2022LS3FF)” and Italian Association for Cancer Research, AIRC (MFAG 2023, Project Code: 28842) to D.L.-B. This work was partially supported by the MUR-PNRR M4C21.3 PE6 project PE00000019 Heal Italia to K.A and by Grants 2019/09679-2 and 2022/06073-9, São Paulo Research Foundation (FAPESP).

## Data and Code Availability

All of the sequencing data used in this paper were obtained from publicly available sources. The main analysis was performed on GSE232278, scRNAseq of BAT db/db and HFD. The bulk RNAseq analysis was performed on GSE112740 BAT HFD and GSE232276 BAT db/db. The FoamSpotteR random forest was trained on GSE97310 and GSE116240, scRNAseq of CD45+ cells of atherosclerotic aorta. GSE116239 bulk RNAseq of foamy and non-foamy macrophages isolated from atherosclerotic aortas via lipid probe-based flow cytometry. The polarization genes were selected using GSE129253, bulk RNAseq of in-vitro polarized macrophages. Validation of MacPolarizeR employed GSE117176, scRNAseq of in-vitro polarized macrophages, and PRJNA626597, scRNAseq of septic adipose tissue. Validation of the role of *Pparg* in macrophages was performed using GSE129253, RNAseq of in vitro downregulated macrophages treated with IL4. Validation of the observation was performed using GSE143437 scRNAseq of Notexin-treated skeletal muscle. The R package code and full tutorials are available on GitHub at https://github.com/andreeedna/MACanalyzeR.

## Authors Contributions

A.N. and F.Z. conceived the study, designed the analyses and interpreted the results. A.N., F.Z., K.A. and D-L.-B. wrote the manuscript and edited the figures. A.N. developed the R package MACanalyzeR. A.N., F.Z. and L.V. performed bioinformatic analyses. F.S. contributed to formal analysis. A.N., F.Z. and L.V. tested MACanalyzeR package. L.S.S., J.C.R.-N. and P.G. performed proteomics analysis. B.Z, J.W.W, S.I., G.R, C.C contributed to the text. D.L.-B. funded the study.

## Conflict of Interests

Authors declare no conflict of interests.

## Materials and Methods

### High Fat Diet Mice

The mice in the study were male 4–6-week-old mice maintained at 22 ± 2°C in a 12-hour light/dark cycle. The mice were fed for 10 weeks with a high-fat diet consisting of a modified AIN-93 diet with an increased lipid content and a reduced carbohydrate content or normal show version (*Table 1*) (Reeves et al., 1993). Two or three mice were kept in each cage and both, water and the diet, were supplied ad libitum. Euthanasia consisted of cervical dislocation followed by decapitation with mice fastened for 16 hours. The Animal Care Committee of the Institute of Biomedical Sciences approved all the experimental protocols (University of São Paulo, Brazil, Protocol CEUA N° 2951040723).

**Table 1.** Diets composition and ingredients (Grams/Kg)

| **Ingredients** | **High Fat Diet** | **Normal Diet** |
| --- | --- | --- |
| *Choline Bitartrate* | 2,6 | 2,5 |
| *Cystine, L* | 3,9 | 1,8 |
| *MIX Vitamínico AIN-93* | 13,8 | 10 |
| *MIX Mineral AIN-93M* | 48 | 35 |
| *Solka Floc* | 74,1 | 50 |
| *Sucrose, Fine Granulated* | 88,9 | 100 |
| *Cornstarch* | 0 | 465 |
| *Lodex 10 (maltodextrin)* | 161,5 | 155 |
| *Casein, Lactic* | 258,4 | 140 |
| *Lard* | 316,6 | 0 |
| *Soybean Oil* | 32,3 | 40 |
| **TOTAL** | 1000 | 999,3 |

### RNAseq Data Analysis

All the RNAseq data were processed using the same pipeline with the same parameters. The bulk RNAseq data were quality-checked using FastQC v0.11.9 and subsequently aligned to the gencode mouse reference genome (mm10) using Hisat2 v2.2.1 (Kim et al., 2019) with default parameters. The number of reads for all RefSeq genes were counted using featureCounts v2.0.3 (Liao et al., 2014) using the multi-mapping option.

The resulting count matrix was analyzed in R using the DESeq2 package v1.40.2 (Love et al., 2014) and Differential Expression was determined using a cutoff significance of padj <0.05. The Gene Set Expression Analysis was made with clusterProfiler v4.8.3. All the heatmaps were plotted using ComplexHeatmap (Wu et al., 2021; Yu et al., 2012).

The linear models were calculated using the *lm* function in R base v4.4.0. The pathway scores were computed on bulk RNAseq data and were used to calculate the linear models.

### Single-cell RNAseq Data Analysis

All the scRNAseq data were processed using the same pipeline with the same parameters. 10x Genomics CellRanger v7.1.0 (Zheng et al., 2017) was used to perform sample demultiplexing, filtering and UMI counting, using the reference transcriptome of *Mus musculus* (mm10) with default parameters. Low quality cells were removed by filtering on the percentage of mitochondrial reads (<10%), number of UMIs (>250 and <30000) and number of detected features (>500).

The scRNAseq analysis of good quality cells was performed in R v4.4.0 using Seurat Package v5.0.3 (Hao et al., 2024). After normalization, we identified the top 2000 variable genes for each library and the libraries were integrated using the CCA method. For all the integrated objects, we performed linear dimensional reduction (PCA), cell clustering and data visualization using UMAP and t-SNE.

Differentially expressed genes that define each cluster were identified using a Wilcoxon Rank Sum test in Seurat and, accompanied by well-known literature gene markers, the DEGs were used to annotate each cell cluster. Bar plots representing cell proportions were plotted using dittoSeq package (Bunis et al., 2021) and the Heatmaps representing gene markers expression were plotted using ComplexHeatmap (Gu, 2022).

Velocity analysis was conducted using two distinct steps. The computation of velocities employed Velocyto v.0.17.17 (La Manno et al., 2018) with BAM files provided as input to generate unspliced and spliced abundance counts in loom format. For the inference of gene-specific RNA velocities, we employed scVelo v0.2.5 to explore cell trajectories. The loom files were processed using a likelihood-based dynamical model, and subsequently, the velocity vectors were projected onto the UMAP projection.

Velocity ranked genes were identified in each cluster using the *scvelo.tl.rank_velocity_genes* function. This function calculates, on velocity expression, the genes that show dynamics that is transcriptomically regulated differently compared to all other clusters.

The velocity ranked genes from the monocytes cluster were used to query the STRING database (Szklarczyk et al., 2021) to obtain a protein-protein interaction network. STRING is a database of protein interactions that are derived from a variety of sources, including experimental data, literature curation, and text mining. The protein-protein interaction network was then used to identify hub genes using the cytoHubba (Chin et al., 2014) plugin for Cytoscape (Shannon et al., 2003). CytoHubba is a collection of algorithms for identifying important nodes in complex networks. The Degrees algorithm was used, which identifies hub genes based on their degree, or the number of connections they have to other genes in the network.

### FoamSpotter Module

FoamSpotteR module is based on a Random Forest (RF) classifier model. RF is an ensemble machine learning technique based on a set of classification trees that are trained on partly independent data subsets. For classification problems, the output is obtained at runtime by majority voting of the trees (Breiman, 2001).

From CD45^+^ cell dataset of atheromatous plaque (GSE), we sub-clustered monocytes and macrophages, resulting in a total of 4976 cells. We annotated these cells as fMAC^+^ and fMAC^-^ based on well-established literature markers, yielding 3914 cells as fMAC^-^ and 1062 cells as fMAC^+^. For model training, we utilized 75% of the cells, while the remaining 25% were reserved as test set. To reduce the dimensionality of the data, we employed the Boruta algorithm (Kursa and Rudnicki, 2010) for feature selection. Boruta is a feature selection algorithm in machine learning that identifies and selects the most important features from a dataset by comparing them to random noise features, helping to improve model performance by focusing on relevant input variables. The top 20 genes were selected based on their importance and subsequently used to train the model (Supplementary Data).

To create and train the RF, we used the R package randomForest (https://CRAN.R-project.org/doc/Rnews/), which implements the original Fortran code in the R environment. The RF was tuned with 500 trees using 4 randomly selected predictors at each split. Furthermore, the FoamSpotteR module provided an additional feature, the FoamDEX Score, which quantifies the probability of a cell developing foam-like features. Values upper 0.5 refer to fMAC. The final RF showed an high level of accuracy, i.e. K = 0.9879.

### MacPolarizeR Module

MacPolarizeR is a module designed to categorize macrophages based on their polarization status. MacPolarizeR operates by clustering cells according to the expression of genes associated in M1 and M2 activation status. To identify genes implicated in the inflammatory process, we interrogated bulk RNAseq of peritoneal c IFN-γ/LPS or IL4 to obtain M1 or M2-like macrophages, respectively (GSE129253). Through a differential gene expression analysis between M1 and M2 cells, we selected the top 75 genes upregulated in pro-inflammatory (M1-like) macrophages and the top 75 genes upregulated in anti-inflammatory (M2-like) macrophages, representing the markers of inflammatory and anti-inflammatory macrophages.

In order to minimize the clustering error, MacPolarizeR was designed as a method based on the k-means algorithm, a point-based clustering that starts with the cluster centers initially placed at arbitrary positions and proceeds by moving at each step the cluster centers (Kawamura et al., 2023). The k-means algorithm was set with 500 iterations and 3 centers corresponding to the classes of macrophages to be clustered: *Inflammatory*, *Transitional* and *Healing*. Additionally, MacPolarizeR calculates an M1 and M2 polarization score for each cell based on the expression of the previously selected genes. The calculated scores can be visualized on a two-dimensional plot, with M1 score on the *X-axis* and M2 score on the *Y-axis* **(Fig. 3A)**. This graphical representation provides a clearer interpretation of macrophage behavior, indicating whether subpopulations lean towards a healing phenotype (second quadrant), a transitional state (first and third quadrants), or an inflammatory state (fourth quadrant). All plots were made by using the ggplot2 package (https://ggplot2.tidyverse.org).

### Pathway Score and PathAnalyzeR Module

To quantify pathway expression in single-cell data, we developed a novel pathway score that represents the weighted average of the expression of genes in the pathways. This calculation can be performed for both individual cells and cell clusters. To allow for comparison across multiple conditions, this calculation considers only normalized gene expressions.

We excluded pathways with at least 5 genes expressed in the dataset and pathways with at least 3 genes expressed in high value (<0.001) in j conditions. For the i-th pathway gene, we first calculated its mean expression level across cell of the j-th condition:

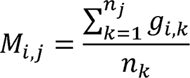

In which n_k_ is the number of cells in the j-th condition, g_i,k_ is the expression level of the i-th gene in the k-th cell in the condition.

The relative expression level of the i-th gene in the j-th condition was then defined as the ratio of M_i,j_ to its average over all cell types.

For condition:

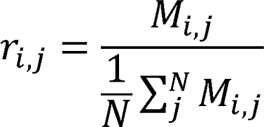

For single cell:

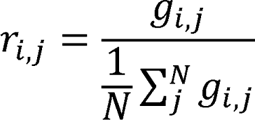

Where r_i,j_ quantifies the relative expression level of gene i in condition j compared to the average expression level of this gene in all conditions.

The weight for each gene is defined as the variance of gene expression:

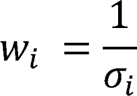

The pathway activity score for the t-th pathway and the j-th condition was then defined as the weighted average of r_i,j_ over all genes included in this pathway:

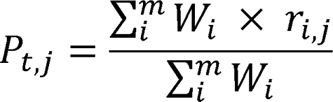

The final score is a positive real number. Scores greater than 1 indicate upregulation of the pathway, while scores less than 1 indicate downregulation of the pathway.

## Notes

https://github.com/andreeedna/MACanalyzeR

### Competing Interest Statement

The authors have declared no competing interest.

